# Pharmacological characterization of FC-12738: A Novel Retro-Inverso Pentapeptide for Treating Neuroinflammation

**DOI:** 10.1101/2023.04.18.537371

**Authors:** Siobhan Ellison

## Abstract

FC-12738, a retro-inverso pentapeptide developed by Neurodegenerative Disease Research, Inc., is currently under investigation for treating neuroinflammation associated with amyotrophic lateral sclerosis (ALS). This study aimed to evaluate the pharmacokinetic properties of FC-12738, including absorption, distribution, metabolism, excretion, and drug-drug interactions. Pharmacokinetics were assessed in Sprague-Dawley rats and beagle dogs following intravenous and subcutaneous administration. Our findings suggest that FC-12738 demonstrates many favorable pharmacokinetic properties, although further optimization may be required to improve CNS penetrance.

## Introduction

Neuroinflammation has been recognized as a key component in the pathogenesis of ALS. In the present study, the pharmacological properties of a novel retro-inverso pentapeptide [FC-12738; (D-Tyr)-(D-Val)-(D-Asp)-(D-Lys)-(D-Arg)], which mirrors the structure of thymopentin (TP5), was evaluated. TP5 is a pentapeptide agonist for the toll-like receptor 2 (TLR2) [1] that constitutes the essential active element of thymopoietin [2,3]. The original description of thymopoietin, as isolated from bovine thymus, reported a 49 amino acid peptide designated Thymopoietin II [4]. In actuality, thymopoietin represents an N-terminal segment of a much larger protein that is normally localized to the nucleus (TMPO; LAP2; www.uniprot.org)[5,6]. The process by which Thymopoietin is generated from its precursor was not revealed by a literature search.

TP5 has been considered as either a primary or adjuvant therapy for treating multiple conditions including, rheumatoid arthritis [7], end-stage renal disease [8], drug-resistant pulmonary tuberculosis [9], cancer [10]. Clinical utilization of TP5 has been hampered by its short half-life *in vivo* [11]. A number of different approaches have been undertaken to try and stabilize the peptide for *in vivo* use, including amino acid modification and fusion to larger carrier proteins [1,12–14]. In the present study, the pharmacological properties of a novel retro-inverso pentapeptide mimetic of TP5 was evaluated. Based on the absorption, distribution, metabolism, and excretion studies, FC-12738 demonstrated rapid absorption and high bioavailability following subcutaneous administration in both Sprague-Dawley rats and beagle dogs. The compound showed low permeability across the Caco-2 cell monolayer and was a poor or non-substrate for efflux transporters. Although detected in brain, the levels were much lower than visceral organs. The low toxicity of FC-12738 is supportive of its further investigation in phase I clinical trials for conditions in which TP5 has been implicated as a possible therapeutic.

## Methods

Absorption studies included permeability and efflux potential evaluation in Caco-2 cells, single-dose pharmacokinetics in Sprague-Dawley (SD) rats, single-dose pharmacokinetics in beagle dogs, single rising dose pharmacokinetics in beagle dogs. Distribution studies included *in vitro* protein binding study, *in vitro* blood to plasma partitioning study, *in vivo* tissue distribution study in SD rats following single subcutaneous injection (SC). Metabolism studies included plasma stability test, hepatocyte stability test, metabolite identification in liver microsomes, metabolite identification in hepatocytes, metabolism in rat plasma, urine, feces and bile. Excretion studies included excretion study in intact SD rats following single SC. Drug-drug interaction studies included Cytochrome P450 (CYP450) induction, inhibition and time-dependent inhibition of FC-12738, and substrate evaluation and inhibition of transporters P-glycoprotein (P-gp), breast cancer resistance protein (BCRP), inhibition of transporters organic anion transporting polypeptide 1B1/3 (OATP1B1/3), organic anion transporter (OAT1/3), organic cation transporter 2 (OCT2), multidrug and toxic compound extrusion 1/2-K (MATE1/2-K). 2.6.4.2

Methods of Tissue and Biospecimen Analysis: The levels of FC-12738 in various tissues, biospecimens, and fluids were measured using a validated LC-MS/MS method. The current method was suitable for the determination of FC-12738 in male beagle dog plasma over the range of 4.00 (lower limit of quantification, LLOQ) to 4000 (upper limit of quantification, ULOQ) ng/mL, using 20 μL of male beagle dog plasma. Ten-fold dilution factor of beagle dog plasma samples had been qualified. The method was suitable for the determination of FC-12738 in male SD rat plasma over the range of 4.00 (LLOQ) to 4000 (ULOQ) ng/mL, using 20 μL of male SD rat plasma. Ten-fold dilution factor of male SD rat plasma samples was qualified. The method was also suitable for the determination of FC-12738 in male SD rat bile over the range of 8.00 (LLOQ) to 8000 (ULOQ) ng/mL and 4.00 (LLOQ) to 4000 (ULOQ) ng/mL in male SD rat feces homogenate and urine, using 40 μL of male SD rat bile, feces homogenate and urine. Samples above the ULOQ could be diluted up to 10-fold or 500-fold using male SD rat bile, feces homogenate and urine. For the determination of FC-12738 in male SD rat brain and kidney homogenate, the peptide was detectable over the range of 4.00 (LLOQ) to 4000 (ULOQ) ng/mL, using 20 μL of male SD rat brain and kidney homogenate. In all cases, the linearity, within-run accuracy and precision, dilution integrity and carryover were successfully evaluated during the qualification and all qualification results met the acceptance criteria.

Assessment of the permeability of FC-12738 across Caco-2 cell monolayers: Caco-2 cells at passage number 50 were seeded on 96-well transport inserts and cultured for 22 days before being used for the transport experiment. FC-12738 was dosed bi-directionally at 2.00, 10.0 and 100 μM with or without 10.0 μM GF120918, a potent inhibitor of efflux transporter(s), such as P-gp and BCRP. Samples were taken at 0 and 120 minutes after incubation and analyzed by LC-MS/MS.

Single-dose pharmacokinetics in SD rats and beagles: Six male SD rats were divided into two groups with 3 animals/group. Animals in Group 1 were administered FC-12738 by single IV administration at 2 mg/kg. Animals in Group 2 were administered FC-12738 by single SC administration at 10 mg/kg. Plasma samples were collected at pre-dose (0), 0.083, 0.25, 0.5, 1, 2, 4, 8 and 24 hours post-dose. Concentrations of FC-12738 in plasma samples were determined by a liquid chromatography tandem mass spectrometry (LC-MS/MS) method. Six non-naïve beagle dogs were divided into two groups with 3 animals/group. Animals in Group 1 were administered FC-12738 by single IV at 1 mg/kg. Animals in Group 2 were administered FC-12738 by single SC at 5 mg/kg. Plasma samples for PK analysis were collected at pre-dose (0), 0.083, 0.25, 0.5, 1, 2, 4, 8 and 24 hours post-dose. Serum samples for antibody microarrays testing were collected at pre-dose (0) and 24 hours post-dose and shipped to sponsor for analysis. Brain samples were collected at 24 hours post-dose and shipped to sponsor for analysis. Concentrations of FC-12738 in plasma samples were determined by LC-MS/MS method.

Single rising dose pharmacokinetics in beagle dogs: Nine non-naïve male beagle dogs were divided into three groups with 3 animals/group. Animals in Groups 1, 2 and 3 were administered FC-12738 by single SC administration at 1, 5 or 25 mg/kg, respectively. Plasma samples were collected at pre-dose (0), 0.083, 0.25, 0.5, 1, 2, 4, 8 and 24 hours post-dose for Groups 1, 2 and 3. Concentrations of FC-12738 in plasma samples were determined by LC-MS/MS method.

*In vitro* protein binding study: The objective of this study was to assess the binding of FC-12738 to CD-1 mouse, SD rat, beagle dog, cynomolgus monkey and human plasma protein by rapid equilibrium dialysis. Plasma containing FC-12738 (0.2, 2 and 10 μM) was dialyzed against phosphate buffer for 6 h in a rapid equilibrium dialysis device at 37 ± 1°C. Warfarin was used as the control compound. Following dialysis, samples were analyzed using LC-MS/MS to determine the relative concentrations in plasma and buffer.

*In vitro* blood to plasma partitioning study: The objective of this study was to determine the blood-to-plasma ratio (K_B/P_) of FC-12738 *in vitro* in whole blood and to estimate the level of partitioning into red blood cells. FC-12738 at 3 concentrations, 0.200, 2.00, and 10.0 μM, was incubated with rat, and dog whole blood at 37°C for 60 min. After incubation, plasma was separated from the incubated blood. The concentrations of analyte in whole blood and plasma were analyzed by LC-MS/MS. Diclofenac (low K_B/P_), chlorthalidone and chloroquine (both high K_B/P_) were used as control compounds to demonstrate the validity of the assay.

Tissue distribution study in SD rats following single subcutaneous injections: Nine male SD rats were divided into three groups with 3 animals/group. Each group was euthanized by CO2 inhalation at 0.5, 2 and 8 hours post-dose, respectively, after receiving a single SC dose of FC-12738 at 10 mg/kg. Blood and eleven types of tissues were collected from each animal at specific time point. Plasma and tissues were processed and then analyzed by LC-MS/MS method.

Plasma stability: The objective of this study was to assess the metabolic stability of FC-12738 in CD-1 mouse, SD rat, beagle dog, cynomolgus monkey and human plasma. FC-12738 at 2.00 μM was incubated with plasma from the above 5 species at 37.0 °C in a water bath for 0, 10, 30, 60 and 120 minutes. Propantheline was used as control compound in CD-1 mouse and human plasma. Enalapril, bisacodyl, and procaine were used as control compounds in SD rat, beagle dog and cynomolgus monkey plasma, respectively. All samples were measured by LC-MS/MS.

Hepatocyte stability: The metabolic stability of FC-12738 in cryopreserved CD-1 mouse, SD rat, beagle dog, cynomolgus monkey, or human hepatocytes was evaluated. FC-12738 was incubated at 1.00 μM with cryopreserved CD-1 mouse, SD rat, beagle dog, cynomolgus monkey, and human hepatocytes at 37.0°C for up to 120 minutes. All samples were measured by LC-MS/MS.

Metabolite identification in liver microsomes: The purpose of this study was to: 1) use LC-UV-HRMS to search for and to identify the metabolites of FC-12738; 2) use LC-MS to determine the relative abundance of FC-12738 and its metabolites; 3) propose the main metabolic pathways of FC-12738. FC-12738, at the concentration of 10 μM, was incubated with liver microsomes at 37 °C for 120 min in the presence of NADPH. The positive control, 7-ethoxycoumarin (7-EC) at 10 μM was run concurrently to assess Phase I metabolic activities in liver microsomes. After incubation, the samples were analyzed by LC-UV-MS. The structures of the metabolites were elucidated based on the characteristics of their MS and MS^2^ data.

Metabolite identification in hepatocytes: The main objectives of this study were to evaluate the metabolism of FC-12738 in mouse, rat, dog, monkey, dog and human hepatocytes. The approaches included: 1) using LC-UV-HRMS to search for and to identify the metabolites of FC-12738; 2) using LC-MS to determine the relative abundance of FC-12738 and its metabolites; 3) propose the main metabolic pathways of FC-12738. FC-12738, at the concentration of 10 μM, was incubated with cryopreserved hepatocytes at 37 °C for 240 min. The positive control, 7-ethoxycoumarin (7-EC) at 30 μM was run concurrently to assess Phase I and Phase II metabolic activities in hepatocytes. After incubation, the samples were analyzed by LC-UV-MS. The structures of the metabolites were elucidated based on the characteristics of their MS and MS^2^ data. The metabolic pathways of FC-12738 in mouse, rat, dog, monkey, dog and human hepatocytes was proposed based on the structures of the metabolites.

Metabolism of FC-12738 in SD rat plasma: The purpose of this study was to evaluate the metabolism of FC-12738 in male SD rat plasma after single IV at the dose of 2 mg/kg and SC at the dose of 10 mg/kg. The approach included: 1) using LC-UV-HRMS to search for and to identify the metabolites of FC-12738; 2) using LC-UV-MS to determine the relative abundance of FC-12738 and its metabolites; 3) propose the formation pathways of major metabolites of FC-12738 in male SD rat plasma. The samples were analyzed by LC-UV-MS. The structures of the metabolite were proposed based on the interpretation of their MS and MS^2^ data, and comparison with reference standards. The metabolic pathways of FC-12738 in male SD rat plasma were proposed based on the structures of the metabolites.

Metabolism of FC-12738 in SD rat urine, feces and bile: The purpose of this study was to evaluate the metabolism of FC-12738 in rat urine, bile and feces after single SC administration at the dose of 10 mg/kg FC-12738. The main approaches were to: 1) use LC-UV-HRMS to search for and to identify the metabolites of FC-12738; 2) use LC-MS to determine the relative abundance of FC-12738 and its metabolites; 3) propose the main metabolic pathways of FC-12738. The samples were analyzed by LC-UV-MS. The structures of the metabolite were proposed based on the interpretation of their MS and MS2 data, and comparison with reference standards. The metabolic pathways of FC-12738 in rat urine, bile and feces were proposed base on the structures of the metabolites.

Excretion study in intact SD rats following single subcutaneous administrations: Six male SD rats were divided into two groups with 3 animals/group. Each rat was administered FC-12738 by single SC administration at 10 mg/kg. Animals in Group 1 were cannulated at the bile duct, and the bile samples were collected from the animals for up to 72 hours post-dose. Animals in Group 2 were put into the metabolic cages, and the feces and urine samples were collected from the animals for up to 168 hours post-dose. Concentrations of FC-12738 in the urine, feces homogenate and bile samples were determined by LC-MS/MS method.

CYP450 inhibition assay: The objective of this study was to determine the potential of FC-12738 to inhibit CYP1A2, CYP2B6, CYP2C8, CYP2C9, CYP2C19, CYP2D6 and CYP3A (with midazolam and testosterone as probe substrate, respectively) in human liver microsomes, using individual probe substrate for each isozyme.

Pooled human liver microsomes were incubated in the presence of FC-12738 (at 0, 0.0300, 0.100, 0.300, 1.00, 3.00, 10.0, 30.0 and 100 μM), NADPH and a selective substrate for each CYP450 isoform. The formation of the selective metabolite from its substrate was measured by liquid chromatography-tandem mass spectrometry.

CYP time-dependent inhibition assay: The objective of this study was to determine the time-dependent inhibitory effects of FC-12738 on the activities of CYP1A2, CYP2B6, CYP2C8, CYP2C9, CYP2C19, CYP2D6 and CYP3A in human liver microsomes using a non-dilution method. Pooled human liver microsomes were pre-incubated for 30 minutes with FC-12738 in the presence or absence of NADPH followed by measurement of remaining enzyme activities.

CYP induction assay: The objective of this study was to evaluate the potential of FC-12738 to induce the mRNAs and activities of CYP450 (CYP) isoforms CYP1A2, CYP2B6, and CYP3A4 in human hepatocytes following *in vitro* administration. Cryopreserved human hepatocytes prepared from three donors were treated with FC-12738 at 0.100, 1.00, 3.00, 10.0 and 15.0 μM at 37°C for a total of 48 hours with the change of incubation medium every 24 hours. The incubation solution after every 24-hour was aspirated for the measurement of Lactate Dehydrogenase (LDH) activity. An aliquot of the incubation solutions from the second dosing was taken at 0, 5 and 24 hours after incubating with the hepatocytes for the measurement of concentration change of FC-12738. After 48 hours incubation, the cells were washed with Hank’s balanced salt solution (HBSS) and incubated with a selective substrate for each CYP450 isoform at 37°C for 30 minutes. The formation of a specific metabolite from its substrate in the incubation solution was quantified by LC-MS/MS method. Cells in the cell culture plate were used to evaluate gene expression by real time Polymerase Chain Reaction (PCR).

Assessment of FC-12738 on the activity of P-gp in the Caco-2 cells: Caco-2 cells at passage number 47 were seeded on 96-well transport inserts and cultured for 23 days before being used for transport experiment. After a 120-minute transport experiment, the bi-directional permeability and efflux ratios of digoxin (10.0 μM), a known P-gp substrate, in the presence of FC-12738 at eight concentrations (0.0457, 0.137, 0.412, 1.23, 3.70, 11.1, 33.3 and 100 μM) or in the absence of FC-12738 were determined and LC-MS/MS was used for bioanalysis. The P-gp activity was represented by the efflux ratios of digoxin in the presence and absence of FC-12738 and the %P-gp Activity values were used to calculate IC_50_.

Assessment of FC-12738 on the activity of BCRP in the Caco-2 cells: Caco-2 cells at passage number 47 were seeded on 96-well transport inserts and cultured for 22 days before being used for the transport experiment. After a 120-min transport experiment, the bi-directional permeability and efflux ratios of estrone 3-sulfate (5.00 μM), a known BCRP substrate, in the presence of FC-12738 at eight concentrations (0.0457, 0.137, 0.412, 1.23, 3.70, 11.1, 33.3 and 100 μM) or in the absence of FC-12738 were determined and LC-MS/MS was used for bioanalysis. The BCRP activity was represented by the efflux ratios of estrone 3 sulfate in the presence and absence of FC-12738 and the %BCRP Activity values were used to calculate IC_50_.

Assessment of FC-12738 on the activity of OATP1B1, OATP1B3, OAT1, OAT3, OCT2, MATE1 and MATE2-K transporters: HEK cells that stably express each transporter (HEK293-OATP1B1, OATP1B3, OAT1, OAT3, OCT2, MATE1, and MATE2-K cells) were incubated with the relevant substrates in the absence and presence of FC-12738. The dosing concentrations of FC-12738 were 0.0457, 0.137, 0.412, 1.23, 3.70, 11.1, 33.3, and 100 μM. After incubation, concentrations of the substrate in samples were analyzed by LC MS/MS. The %VC (%Vehicle Control, the percentage of the mean transport activity values in the presence and absence of test compound) values were used for half maximal inhibitory concentration (IC_50_) calculation.

Comparison of TP5 and FC-12738 in plasma and brain in male CD-1 mice after a single IP dose: Three animals were dosed via injection in the lower right quadrant of the abdomen for intraperitoneal injections at 10 mg/kg (10 mL/kg). The peptides were suspended in solutions consisting of 10% N,N-Dimethlyacetamide (DMA), 20% propylene glycol (PG), 70% H_2_O. All blood samples (30-50 μL per sample) were taken via appropriate vein (saphenous or submandibular vein) at 0.5, 1, 2, 4, and 8 h for the brain tissue distribution group. Blood samples were collected in Greiner MiniCollect K2EDTA tubes, placed on ice, and within 30 minutes, centrifuged at 15,000g for 5 min to obtain plasma samples. All plasma samples were stored at –70C until analysis. The whole brain was harvested at selected time points, immediately rinsed in water briefly, and blotted dry with a paper towel. The whole brain was weighed and three volumes of PBS buffer (pH 7.4) was added to one volume of each tissue sample which was homogenized by a tissue homogenizer until fine tissue particles were completely dispersed or emulsified. The tissue homogenate samples were stored at −70 °C until analysis.

Plasma samples were prepared as follows. Three volumes of acetonitrile containing internal standard was added to one volume of plasma to precipitate proteins. Samples were centrifuged (3000 ×g for 10 min) and supernatant was removed for analysis by LCMS/MS. Calibration standards and quality controls were made by preparation of a 1 mg/mL stock solution and subsequently a series of working solutions in methanol : water (1/:1, v/v) which were spiked into blank plasma to yield a series of calibration standard samples in the range of 1 ng/mL to 10 μg/mL and quality control samples at three concentration levels (low, middle and high). All incurred PK/PD plasma samples were treated identically to the calibration standards and quality control samples. LC-MS/MS analysis was performed utilizing multiple reaction monitoring for detection of characteristic ions for each drug candidate, additional related analytes and internal standard.

Tissue samples were prepared as follows. Three volumes of PBS buffer (pH 7.4) was added to one volume of each tissue sample which was then homogenized to obtain each tissue homogenate sample. Subsequently, three volumes of acetonitrile containing internal standard was added to one volume of each tissue homogenate, and the mixture was vortexed, centrifuged (3000 g for 10 min) and supernatant was removed for analysis by LC-MS/MS. Calibration standards were made by preparation of a 1 mg/mL stock solution and subsequently a series of working solutions in methanol : water (1/:1, v/v) which were spiked into blank tissue homogenate to yield a series of calibration standard samples in the range of 1 ng/mL to 10 μg/mL. All incurred PK/PD tissue samples were treated identically to the calibration standards. LC-MS/MS analysis was performed utilizing multiple reaction monitoring for detection of characteristic ions for each drug candidate, additional related analytes and internal standard.

## Results and Discussion

### Caco cell permeability

Caco cells were exposed to increasing concentrations of FC-12738 in the presence or absence of GF120918, a potent inhibitor of efflux transporter(s), such as P-gp and BCRP. In the absence of GF120918, at 2.00, 10.0 and 100 μM dosing concentrations, FC-12738 showed the mean apparent permeability coefficient (Papp) values of <0.241, 0.107 and 0.0949 × 10-6 cm/s, respectively, in the apical to basolateral (A to B) direction, and <0.217, 0.0946 and 0.117 × 10-6 cm/s, respectively, in the B to A direction. Therefore, the efflux ratio (ERa) couldn’t be calculated at the dosing concentration of 2.00 μM. At the dosing concentrations of 10.0 and 100 μM, no efflux was observed as evidenced by the ERa values of 0.880 and 1.24, respectively. In the presence of GF120918, FC-12738 showed the mean Papp values of <0.237, 0.0900 and 0.0752 × 10-6 cm/s, respectively, at the three tested dosing concentrations, in the A to B direction, and <0.213, 0.101 and 0.0927 × 10-6 cm/s, respectively, in the B to A direction. Therefore, the efflux ratio (ERi) couldn’t be calculated at the dosing concentration of 2.00 μM, and the ERi values were 1.13 and 1.23, respectively, at the dosing concentrations of 10.0 and 100 μM (**Table 1**). In conclusion, under the conditions tested, FC-12738 demonstrated low permeable across the Caco-2 cell monolayer and was a poor or non-substrate for efflux transporters.

**Table 1.** Summary of the bidirectional permeability of FC-12738 in the Caco-2 cells.

| Conc. (μM) | + / - GF120918 | Mean P<sub>app</sub><br>($\times 10^{-6}$ cm/s) | | Efflux Ratio | Mean %Solution Recovery | |
| --- | --- | --- | --- | --- | --- | --- |
|  |  | A to B | B to A |  | A to B | B to A |
| 2.00 | - | <0.241 | <0.217 | NA | <94.1 | <99.0 |
|  | + | <0.237 | <0.213 | NA | <92.9 | <96.5 |
| 10.0 | - | 0.107 | 0.0946 | 0.880 | 90.2 | 100.5 |
|  | + | 0.0900 | 0.101 | 1.13 | 89.9 | 96.5 |
| 100 | - | 0.0949 | 0.117 | 1.24 | 89.1 | 95.3 |
|  | + | 0.0752 | 0.0927 | 1.23 | 86.9 | 97.0 |

### Pharmacokinetics

After IV administration of FC-12738 at 2 mg/kg in male SD rats, FC-12738 showed a plasma clearance (Cl) of

11.8±0.569 mL/min/kg, half-life (T1/2) at 0.318±0.0105 h. The volume of distribution (Vdss) was 0.309±0.0165 L/kg, the area under the plasma concentration time curve from time zero to the last quantifiable concentration (AUC0-last) value was 2800±131 ng·h/mL. Following SC administration of FC-12738 at 10 mg/kg in male SD rats, AUC0-last value of FC-12738 was 13000±379 ng·h/mL. The Cmax value of FC-12738 was 8430±637 ng/mL, while Cmax was reached at 0.417±0.144 h. The bioavailability was 91.9% at 10 mg/kg. All animals had tolerated the FC-12738 at the dosing levels during the entire course of the study. No adverse effect was observed during the in-life phase of the study. The mean PK parameters of FC-12738 in male SD rats are shown in **Table 2**.

**Table 2.**
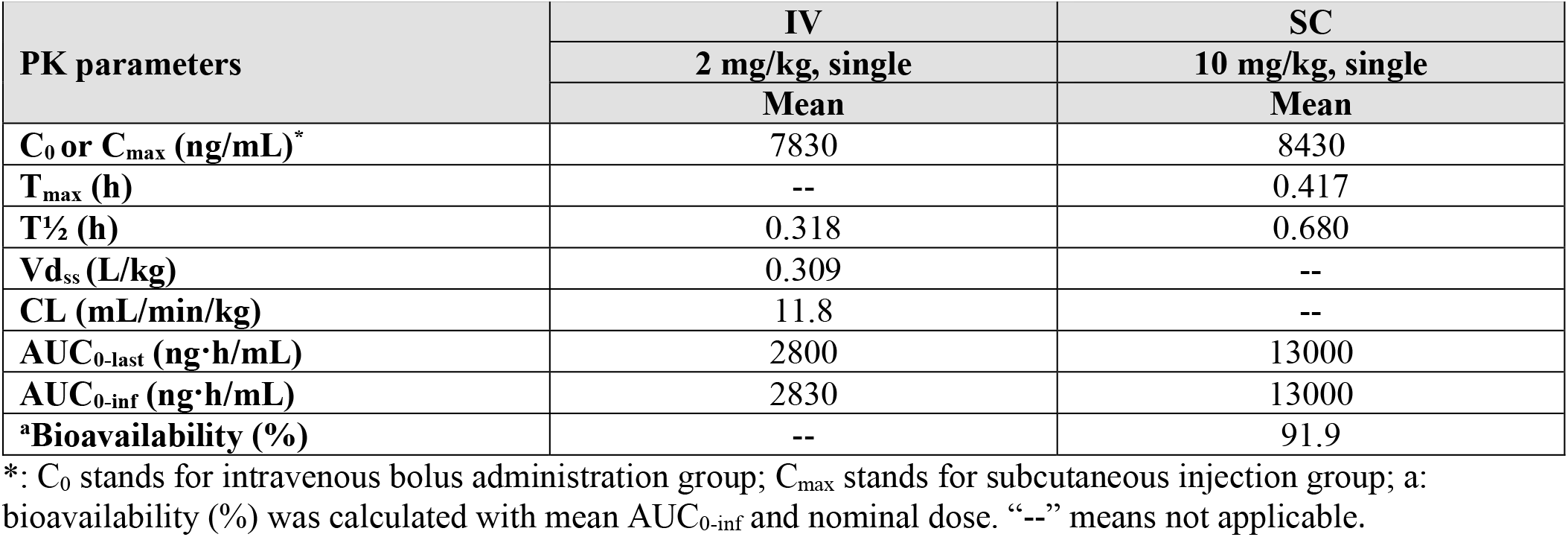
Mean pharmacokinetic parameters of FC-12738 in male SD rats.

| PK parameters | IV | SC |
| --- | --- | --- |
|  | 2 mg/kg, single | 10 mg/kg, single |
|  | Mean | Mean |
| C<sub>0</sub> or C<sub>max</sub> (ng/mL)* | 7830 | 8430 |
| T<sub>max</sub> (h) | -- | 0.417 |
| T<sub>1/2</sub> (h) | 0.318 | 0.680 |
| Vd<sub>ss</sub> (L/kg) | 0.309 | -- |
| CL (mL/min/kg) | 11.8 | -- |
| AUC<sub>0-last</sub> (ng·h/mL) | 2800 | 13000 |
| AUC<sub>0-inf</sub> (ng·h/mL) | 2830 | 13000 |
| <sup>a</sup>Bioavailability (%) | -- | 91.9 |

\*: C<sub>0</sub> stands for intravenous bolus administration group; C<sub>max</sub> stands for subcutaneous injection group; a: bioavailability (%) was calculated with mean AUC<sub>0-inf</sub> and nominal dose. "--" means not applicable.

After IV administration of FC-12738 at 1 mg/kg in male beagle dogs, FC-12738 showed Cl of 5.85±0.940 mL/min/kg, T1/2 at 0.680±0.0504 h **(Table 3**). Vdss was 0.302±0.0636 L/kg, the area under the plasma concentration time curve from time zero to AUC0-last value was 2850±439 ng·h/mL. Following single SC administration of FC-12738 at 5 mg/kg in male beagle dogs, AUC0-last value of FC-12738 was 16100±2510 ng·h/mL, the Cmax value was 7300±1050 ng/mL, while Tmax was reached at 0.833±0.289 h. The SC bioavailability was 112% at 5 mg/kg. All animals tolerated FC-12738 at the dosing levels during the entire course of the study. No adverse effect was observed during the in-life phase of the study.

**Table 3.** The mean plasma concentrations (ng/mL), tissue concentrations (ng/g) and pharmacokinetic parameters of FC-12738 in male SD rats (n=3 per time point)

| Time (h)<br>Matrix | Concentration (ng/mL or ng/g) |  |  | Parameters |  |  |
| --- | --- | --- | --- | --- | --- | --- |
| | 0.5 | 2 | 8 | $C_{max}$<br>(ng/mL or<br>ng/g) | $T_{max}$<br>(h) | $AUC_{0-last}$<br>(ng·h/mL or<br>ng·h/g) |
| Plasma | 14200 | 1790 | 5.04 | 14200 | 0.500 | 14400 |
| Liver | 1270 | 616 | 269 | 1270 | 0.500 | 4190 |
| Kidney | 28500 | 17300 | 14300 | 28500 | 0.500 | 135000 |
| Stomach | 3370 | 555 | 96.7 | 3370 | 0.500 | 4760 |
| Lung | 5330 | 752 | 109 | 5330 | 0.500 | 6840 |
| Small Intestine | 2060 | 775 | 188 | 2060 | 0.500 | 4970 |
| Large Intestine | 2150 | 430 | 226 | 2150 | 0.500 | 4040 |
| Heart | 2150 | 365 | ND | ND | ND | ND |
| Muscle | 1490 | 191 | 46.4 | 1490 | 0.500 | 1930 |
| Spleen | 1490 | 345 | 239 | 1490 | 0.500 | 3280 |
| Fat | 1070 | 103 | ND | ND | ND | ND |
| Brain | 128 | ND | ND | ND | ND | ND |

ND: not determined as more than half (>50%) of the individual values were not quantifiable

**Table 1.** Mean pharmacokinetic parameters of FC-12738 in male beagle dogs.

| PK Parameters | IV | SC |
| --- | --- | --- |
|  | 1 mg/kg, single | 5 mg/kg, single |
|  | Mean | Mean |
| $C_0$ or $C_{max}$ (ng/mL)* | 4820 | 7300 |
| $t_{max}$ (h) | -- | 0.833 |
| $t_{1/2}$ (h) | 0.680 | 0.884 |
| $V_{dss}$ (L/kg) | 0.302 | -- |
| CL (mL/min/kg) | 5.85 | -- |
| AUC<sub>0-last</sub> (ng·h/mL) | 2850 | 16100 |
| AUC<sub>0-inf</sub> (ng·h/mL) | 2900 | 16200 |
| Bioavailability (%)<sup>a</sup> | -- | 112 |

\*: $C_0$ stands for intravenous bolus administration group; $C_{max}$ stands for subcutaneous injection group.
a: Bioavailability (%) was calculated with mean AUC<sub>0-last</sub> and nominal dose. "--" means not applicable.

Following SC administration of FC-12738 at 1, 5 and 25 mg/kg in male beagle dogs, the area under the plasma concentration time curve from time zero to AUC0-last values of FC-12738 were 2760±120, 16700±3260 and 72800±4920 ng·h/mL, respectively. The Cmax values were 925±39.7, 7090±935 and 33300±4070 ng/mL, while Tmax was reached at 1.00, 1.00 and 0.667±0.289 h, respectively. The systemic exposure (AUC0-last and Cmax) of FC-12738 increased with dose proportionally as dose levels increased from 1 mg/kg to 5 mg/kg, 5 mg/kg to 25 mg/kg and 1 mg/kg to 25 mg/kg in male beagle dogs. The mean PK parameters of FC-12738 are shown in **Table 4**.

**Table 2.** Mean pharmacokinetic parameters of FC-12738 in male beagle dogs.

| PK parameters | SC |  |  |
| --- | --- | --- | --- |
|  | 1 mg/kg, single | 5 mg/kg, single | 25 mg/kg, single |
|  | Mean | Mean | Mean |
| $C_{max}$ (ng/mL)* | 925 | 7090 | 33300 |
| $T_{max}$ (h) | 1.00 | 1.00 | 0.667 |
| $T_{1/2}$ (h) | 0.957 | 0.906 | 2.01 |
| AUC<sub>0-last</sub> (ng·h/mL) | 2760 | 16700 | 72800 |
| AUC<sub>0-inf</sub> (ng·h/mL) | 2770 | 16800 | 72800 |

\*: $C_{max}$ stands for subcutaneous injection group

**Table 4.** The relative abundance of metabolites and FC-12738 after incubated with human liver microsomes.

| Metabolite Code | $[M+H]^+$<br>$m/z$ | RT (min) | Relative Abundance (MS Peak Area %Total) <sup>a</sup> | Metabolic Pathways |
| --- | --- | --- | --- | --- |
|  |  |  | human |  |
| FC-12738 | 680.3721 | 21.47 | 100.00 | NA, Parent Drug |

RT: Retention time of LC-MS; NA: Not applicable. <sup>a</sup>

### Protein binding Study

At 0.2, 2 and 10 μM, the binding of FC-12738 was 55.1%, 54.5% and 52.1% in CD-1 mouse plasma, 51.8%, 55.7% and 50.7% in SD rat plasma, 50.8%, 53.9% and 30.9% in beagle dog plasma, 46.2%, 46.1% and 26.4% in cynomolgus monkey plasma, 57.2%, 53.0% and 32.7% in human plasma, respectively. The recovery of FC-12738 from dialysis wells was acceptable, ranging from 100.9% to 120.3%, indicating that the compound was stable during the dialysis process. In conclusion, FC-12738 exhibited a moderate binding to plasma proteins in CD-1 mouse and SD rat, a low to moderate binding to plasma proteins in beagle dog and human plasma, and a low binding to plasma proteins in cynomolgus monkey, at all three test concentrations (0.2, 2 and 10 μM).

### Blood to plasma partitioning study

At 0.200, 2.00, and 10.0 μM, the mean KB/P values of FC-12738 were 0.48, 0.50, and 0.47 in rat blood, 0.57, 0.55, and 0.58 in dog blood whole blood, respectively. The mean erythrocyte to plasma ratio (KE/P) values were <0.21, <0.11, and <0.03 in rat blood, <0.18, 0.07, and <0.14 in dog blood whole blood, respectively. Following a 60-min incubation, the %Recovery of FC-12738 was acceptable, ranging from 89.98 to 101.03. In conclusion, FC-12738 exhibited a low partitioning into erythrocytes of rat and dog. No significant species difference or concentration-dependent partitioning was observed.

### Tissue distribution study in SD rats following single subcutaneous injections

After single SC administration of FC-12738 at 10 mg/kg to male SD rats, FC-12738 was distributed to all tissues within 0.5 hour post-dose. C_max_ were observed at 0.5 hour post-dose in most of tissues. The highest tissue exposure (AUC_0-last_) was observed in kidney (135000 ng·h/g) and lung (6840 ng·h/g), followed by small intestine, stomach, liver, large intestine, spleen and muscle. The heart, fat and brain exposures (AUC_0-last_) were not determined due to most concentrations were below the LLOQ (40 ng/g). All animals had tolerated the compound at the dosing levels during the entire course of the study. No adverse effect was observed during the in-life phase of the study. The mean plasma concentrations (ng/mL), tissue concentrations (ng/g) and pharmacokinetic parameters of FC-12738 in male SD rats are shown in **Table 5**.

**Table 5.** The metabolic changes and relative abundance of FC-12738 and its detected metabolites in hepatocytes.

| Metabolite Code | [M+H]<sup>+</sup><br>m/z | RT<br>(min) | Relative Abundance (UV%)<sup>a</sup> |  |  |  |  | Metabolic Pathways |
| --- | --- | --- | --- | --- | --- | --- | --- | --- |
|  |  |  | Mouse | Rat | Dog | Monkey | human |  |
| *M1 | 842.4268 | 20.24 | 0.33 | 0.23 | 0.18 | 0.27 | 0.21 | Glucose conjugation (P + C<sub>6</sub>H<sub>10</sub>O<sub>5</sub>) |
| *M2 | 842.4252 | 21.07 | 3.76 | 2.87 | 2.85 | 2.60 | 2.66 | Glucose conjugation (P + C<sub>6</sub>H<sub>10</sub>O<sub>5</sub>) |
| FC-12738 | 680.3719 | 21.22 | 88.92 | 87.56 | 80.76 | 89.24 | 91.82 | NA |
| *M3 | 842.4255 | 22.63 | 5.15 | 3.18 | 4.17 | 3.14 | 3.28 | Glucose conjugation (P + C<sub>6</sub>H<sub>10</sub>O<sub>5</sub>) |
| *M4 | 842.4258 | 24.82 | 1.84 | 1.31 | 1.48 | 1.19 | 1.28 | Glucose conjugation (P + C<sub>6</sub>H<sub>10</sub>O<sub>5</sub>) |

P: parent drug; NA: Not applicable; \*: metabolites were detected in NC (without hepatocytes);

### Plasma stability

At different time points 0, 10, 30, 60, and 120 minutes, FC-12738 showed %Remaining values of 100.0, 103.6, 100.4, 110.6 and 100.1 in CD-1 mouse plasma; 100.0, 104.4, 102.2, 112.9 and 103.6 in SD rat plasma; 100.0, 106.2, 105.1, 105.8 and 100.6 in beagle dog plasma; 100.0, 104.8, 104.9, 104.7 and 102.5 in cynomolgus monkey plasma; 100.0, 93.3, 93.6, 90.6, and 85.6 in human plasma, respectively. FC-12738 has a half-life of >372.7 minutes in CD-1 mouse, SD rat, beagle dog, cynomolgus monkey and human plasma, respectively. In conclusion, FC-12738 was stable in CD-1 mouse, SD rat, beagle dog, cynomolgus monkey and human plasma after 120 minutes incubation.

### Hepatocyte stability

The half-life values of FC-12738 were >372.7, >372.7, >372.7 >372.7 and >372.7 (min) after 120 minutes incubations, corresponding to the intrinsic clearance CL_int(hep)_ (μL/min/10^6^ cells) values of <1.9, <1.9, <1.9, <1.9, and <1.9; and hepatic clearance CL_int(liver)_ (mL/min/kg) values of <22.1, <8.7, <12.8, <6.7 and <5.2 in mouse, rat, dog, monkey, and human hepatocytes, respectively. These results indicated that FC-12738 was slowly metabolized in CD-1 mouse, SD rat, beagle dog, cynomolgus monkey and human hepatocytes.

### Metabolite identification in liver microsomes

No metabolites of FC-12738 were detected and identified from human liver microsomes. Based on the identified metabolites, FC-12738 (MW = 679.4) was the major component in human liver microsomes after incubation at 37 °C for 120 min. The relative abundances (UV%) of parent drug and its metabolites in human liver microsomal incubations are shown in **Table 6**.

**Table 6.** The relative MS abundance of FC-12738 and its metabolites in rat plasma.

| Metabolite Code | [M+2H]<sup>2+</sup> | RT<br>(min) | Male |  | Metabolic Pathways |
| --- | --- | --- | --- | --- | --- |
|  |  |  | 0-2 h (IV) | 0-4 h (SC) |  |
| M2 | *421.7159 | 19.87 | 0.66 | 1.43 | Glucose conjugation (P + C<sub>6</sub>H<sub>10</sub>O<sub>5</sub>) |
| FC-12738 | *340.6898 | 19.81 | 97.10 | 96.57 | NA |
| M3 | *421.7160 | 22.46 | 2.03 | 1.74 | Glucose conjugation (P + C<sub>6</sub>H<sub>10</sub>O<sub>5</sub>) |
| M4 | *421.7158 | 24.56 | 0.20 | 0.26 | Glucose conjugation (P + C<sub>6</sub>H<sub>10</sub>O<sub>5</sub>) |

NA: not applicable. RT: Retention time of LC-MS; NA: Not applicable; P: Parent; \*: value of [M+2H]<sup>2+</sup>.

### Metabolite identification in hepatocytes

A total of 4 metabolites of FC-12738 were detected and identified from mouse, rat, dog, monkey, and human hepatocytes. The metabolites were assigned as M1-M4: glucose conjugation metabolites (MW = 841.92, P + C6H10O5). Based on the identified metabolites, the major metabolic pathway of FC-12738 in hepatocytes was proposed as glucose conjugation. The metabolites formed in incubation of human hepatocytes were detected in hepatocytes of at least one animal species. The relative abundance (UV%) of FC-12738 and each putative metabolites were provided in **Table 7**. The proposed major metabolic pathway in hepatocytes of FC-12738 was shown in **Error! Reference source not found**..

**Table 7.** The relative MS abundance of FC-12738 and its metabolites in SD rat urine, bile and feces.

| Metabolite Code | [M+2H]<sup>2+</sup> | RT (min) | Male |  |  | Metabolic Pathways |
| --- | --- | --- | --- | --- | --- | --- |
|  |  |  | Urine | Feces | Bile |  |
| FC-12738 | *340.6896 | 20.03 | 99.93 | 100.00 | 100.00 | NA |
| M3 | *421.7161 | 22.49 | 0.06 | ND | ND | Glucose conjugation (P + C<sub>6</sub>H<sub>10</sub>O<sub>5</sub>) |
| M4 | *421.7153 | 24.66 | 0.01 | ND | ND | Glucose conjugation (P + C<sub>6</sub>H<sub>10</sub>O<sub>5</sub>) |

NA: not applicable. RT: Retention time of LC-MS; NA: Not applicable; P: Parent; \*: value of [M+2H]<sup>2+</sup>.

### Metabolism of FC-12738 in SD rat plasma

In addition to unchanged FC-12738 (MW = 679.37), a total of 3 metabolites of FC-12738 were detected and identified in male SD rat plasma. The metabolites were assigned as: M2-M4: glucose conjugation metabolites (MW = 841.92, P + C_6_H_10_O_5_). The relative abundance of FC-12738 and its metabolites is shown in **Table 8**. Based on the identified metabolites, the proposed formation pathway of major metabolites of FC-12738 in male SD rat plasma was glucose conjugation. The proposed formation pathways of major metabolites of FC-12738 in SD rat plasma are shown in Error! Reference source not found..

**Table 8.** Accumulative Dose Recovery (%) of FC-12738 in Bile, Urine and Feces of Male SD rats Following Single subcutaneous injection administration of FC-12738 at 10 mg/kg.

| Matrix | Collection Time (h) | Male |  |
| --- | --- | --- | --- |
|  |  | Mean | SD |
| Urine | 0 | 0.00 | -- |
|  | 4 | 56.2 | 26.2 |
|  | 8 | 78.2 | 8.68 |
|  | 24 | 84.0 | 6.22 |
|  | 48 | 86.1 | 6.51 |
|  | 72 | 86.4 | 6.35 |
|  | 96 | 86.4 | 6.31 |
|  | 120 | 86.6 | 6.25 |
|  | 144 | 86.7 | 6.19 |
|  | 168 | 86.7 | 6.25 |
| Feces | 0 | 0.00 | -- |
|  | 4 | 0.00168 | 0.00292 |
|  | 8 | 0.00239 | 0.00414 |
|  | 24 | 1.74 | 0.962 |
|  | 48 | 2.08 | 1.20 |
|  | 72 | 2.14 | 1.21 |
|  | 96 | 2.16 | 1.22 |
|  | 120 | 2.18 | 1.22 |
|  | 144 | 2.19 | 1.23 |
|  |  | Mean | SD |
|  | 168 | 2.19 | 1.22 |
| Total | 0 | 0.00 | -- |
|  | 4 | 56.2 | 26.2 |
|  | 8 | 78.2 | 8.68 |
|  | 24 | 85.8 | 5.85 |
|  | 48 | 88.2 | 6.00 |
|  | 72 | 88.5 | 5.81 |
|  | 96 | 88.6 | 5.76 |
|  | 120 | 88.7 | 5.71 |
|  | 144 | 88.9 | 5.71 |
|  | 168 | 88.9 | 5.76 |
| Bile | 0 | 0.00 | -- |
|  | 4 | 0.284 | 0.279 |
|  | 12 | 0.314 | 0.270 |
|  | 24 | 0.324 | 0.271 |
|  | 48 | 0.331 | 0.264 |
|  | 72 | 0.336 | 0.260 |

### Metabolism of FC-12738 in SD rat urine, feces and bile

A total of 2 metabolites of FC-12738 were detected and identified in rat urine, bile and feces. The metabolites were assigned as: M3 and M4: glucose conjugation metabolites (MW = 841.92, P + C_6_H_10_O_5_). The relative abundance of FC-12738 and its metabolites shown as in **Table 9**. Based on the identified metabolites, the major metabolic pathway of FC-12738 in rat urine, bile and feces was proposed as glucose conjugation. The proposed major metabolic pathways of FC-12738 in rat urine, bile and feces are presented in **Figure 1**.

**Figure 1.**
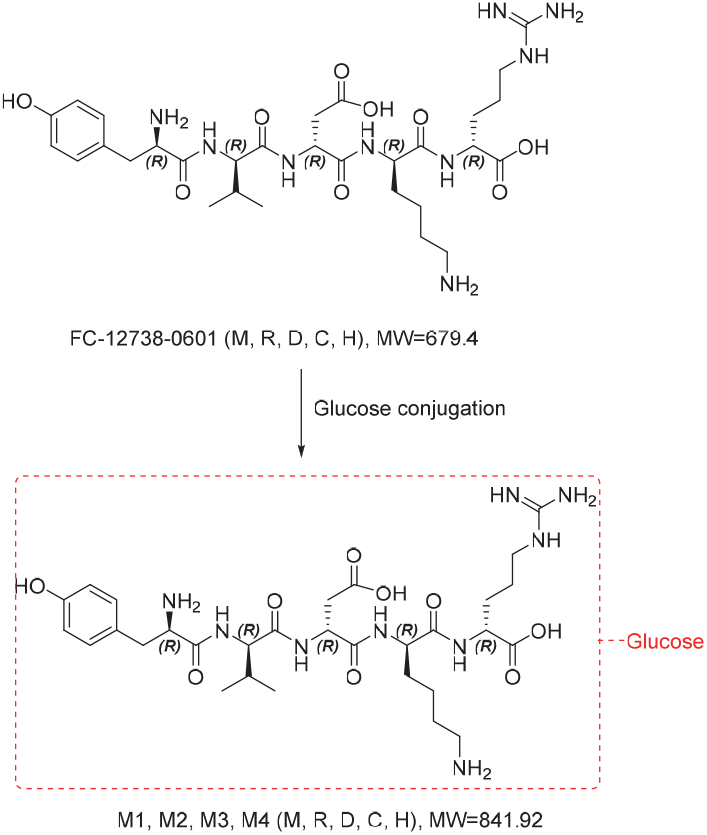
Proposed metabolic pathway of FC-12738 in hepatocytes, plasma, urine, bile, and feces.

### Excretion study in intact SD rats following single subcutaneous administrations

Following a single SC administration of FC-12738 at 10 mg/kg, the mean drug recovery of FC-12738 from urine and feces in the male intact rats at 168 hours post-dose were 86.7±6.25% and 2.19±1.22%, respectively, of the nominal dose. The total recovery rate was 88.9±5.76% of the dosage, and urine was the main excretion route. Drug recovery of FC-12738 from bile in the male bile-duct-cannulated (BDC) rats at 72 hours post-dose was 0.336±0.260% of the nominal dose. All animals had tolerated the compound at the dosing levels during the entire course of the study. No adverse effect was observed during the in-life phase of the study. The accumulative dose recovery (%) of FC-12738 in bile, urine and feces of male SD rats following single SC administration of FC-12738 at 10 mg/kg is shown in **Table**.

### CYP450 inhibition assay

The remaining activities of CYP1A2, CYP2B6, CYP2C8, CYP2C9, CYP2C19, CYP2D6 and CYP3A (with midazolam and testosterone as probe substrate, respectively) treated with FC-12738 were no less than 97.4%, 99.9%, 99.0%, 96.7%, 95.9%, 95.1%, 95.1% and 97.2%, respectively. Therefore, the IC_50_ values of FC-12738 on CYP1A2, CYP2B6, CYP2C8, CYP2C9, CYP2C19, CYP2D6 and CYP3A (with midazolam and testosterone as probe substrate, respectively) were all greater than 100 μM.

### CYP time-dependent inhibition assay

The fold of IC_50_ shift for CYP1A2, CYP2B6, CYP2C8, CYP2C9, CYP2C19, CYP2D6, CYP3A (midazolam as substrate) and CYP3A (testosterone as substrate) could not be calculated because the IC50 values were greater than 100 μM for the pre-incubation with or without NADPH. Using an IC50 shift greater than or equal to 1.5-fold as the cutoff for time-dependent inactivation, it was concluded that FC-12738 showed no time-dependent inhibition on any of the CYP isozymes tested.

### CYP induction assay

The released LDH data show that FC-12738 at 0.100, 1.00, 3.00, 10.0 and 15.0 μM has no cytotoxicity in all three donors. The data from enzyme activities levels and gene expression levels in all three donors show that FC-12738 at the concentrations ranging from 0.100 to 15.0 μM is not considered as an inducer for CYP1A2, CYP2B6 and CYP3A4. The metabolic stability data show that following incubation with human hepatocytes for 24 hours, the % remaining for FC-12738 at concentrations tested were found to be 107.4, 105.0, 47.1, 69.1, 86.4 in Donor 1, 89.2, 112.4, 61.9, 68.7, 89.3 in Donor 2, and 93.8, 100.3, 55.0, 61.0, 91.7 in Donor 3, respectively.

### Assessment of FC-12738 on the activity of P-gp and BCRP in the Caco-2 cells

Under the conditions tested, the %P gp Activity values of Caco-2 cells treated with FC-12738 were no less than 93.9, therefore, the IC_50_ value of FC-12738 on P-gp was greater than 100 μM. Under the conditions tested, the %BCRP Activity values of Caco-2 cells treated with FC-12738 were no less than 93.3, therefore, the IC_50_ value of FC-12738 on BCRP was greater than 100 μM.

### Assessment of FC-12738 on the activity of OATP1B1, OATP1B3, OAT1, OAT3, OCT2, MATE1 and MATE2-K transporters

The %VC values of HEK293-OATP1B1, OATP1B3, OAT1, OAT3, OCT2, MATE1, and MATE2-K cells treated with FC-12738 were no less than 99.4, 89.6, 98.4, 98.8, 99.6, 91.3, and 62.6, respectively. Therefore, the IC_50_ values of FC-12738 on OATP1B1, OATP1B3, OAT1, OAT3, OCT2, MATE1, and MATE2-K were greater than 100 μM.

### Comparison of TP5 and FC-12738 levels in plasma and brain in male CD-1 mice after a single IP dose

The plasma and brain levels of TP5 and FC-12738 were compared in male CD-1 mice after an IP injection of each peptide at a concentration of 10mg/ml. Vehicle formulations for the two peptides were identical. In the first 2 hours after injection, the levels of FC-12738 in plasma and brain were substantially higher than that of TP5 (Table 11). These findings are consistent with the idea that FC-12738 has a substantially longer half-life in plasma than TP5, allowing for greater uptake in brain.

**Table 11.** Levels of TP5 and FC-12738 in plasma and brain of CD-1 mice.

| Analyte | Matrix | Time (h) | Plasma or Tissue Homogenate Conc (ng/ml) n=3 |  |  | Mean (ng/ml) | Mean Tissue (ng/g) |
| --- | --- | --- | --- | --- | --- | --- | --- |
| FC-12738 | Plasma | 0.5 | 2230 | 1790 | 2260 | **2093** | N/A |
|  |  | 1 | 885 | 1320 | 1420 | **1208** |  |
|  |  | 2 | 543 | 592 | 745 | **627** |  |
|  |  | 4 | 97.9 | 74.6 | 79.8 | **84.1** |  |
|  |  | 8 | 19.1 | 17.5 | 13.4 | **16.7** |  |
|  | Brain | 0.5 | 17.4 | 17.6 | 14.1 | **16.4** | **65.5** |
|  |  | 1 | 8.28 | 10.1 | 8.41 | **8.93** | **35.7** |
|  |  | 2 | 6.96 | 5.73 | 7.47 | **6.72** | **26.9** |
|  |  | 4 | 1.76 | 1.22 | 1.06 | **1.35** | **5.39** |
|  |  | 8 | ND | ND | ND | **ND** | **ND** |
| TP5 | Plasma | 0.5 | 144 | 136 | 115 | **132** | N/A |
|  |  | 1 | 80.1 | 107 | 85.7 | **90.9** |  |
|  |  | 2 | 56.5 | 67.2 | 54.8 | **59.5** |  |
|  |  | 4 | 53.4 | 42.2 | 51.3 | **49** |  |
|  |  | 8 | 28.7 | 30.6 | 23.6 | **27.6** |  |
|  | Brain | 0.5 | 2.34 | 2.17 | 3.18 | **2.56** | **10.3** |
|  |  | 1 | 1.69 | 1.78 | 2.20 | **1.89** | **7.56** |
|  |  | 2 | 1.64 | 1.28 | 1.22 | **1.38** | **5.52** |
|  |  | 4 | 1.16 | 1.03 | 1.08 | **1.09** | **4.36** |
|  |  | 8 | ND | ND | ND | **ND** | **ND** |

## Conclusions

In regard to absorption, FC-12738 showed low permeability across the Caco-2 cell monolayer and was a poor or non-substrate for efflux transporters. Following single SC administration of FC-12738 at 10 mg/kg in male SD rats and at 5 mg/kg in male beagle dogs, FC-12738 showed great bioavailability. And in single rising dose study in beagle dogs at 1, 5 and 25 mg/kg, the systemic exposure (AUC_0-last_ and C_max_) of FC-12738 increased with dose proportionally as dose levels increased from 1 mg/kg to 5 mg/kg, 5 mg/kg to 25 mg/kg and 1 mg/kg to 25 mg/kg in male beagle dogs.

In regard to distribution in plasma, FC-12738 exhibited a moderate binding to plasma proteins in CD-1 mouse and SD rat plasma, a low to moderate binding to plasma proteins in beagle dog and human plasma, and a low binding to plasma proteins in cynomolgus monkey, at all three test concentrations tested (0.2, 2 and 10 μM). FC-12738 exhibited a low partitioning into erythrocytes of rat and dog. No significant species difference or concentration-dependent partitioning was observed.

In the tissue distribution study in SD rats following single SC, the highest tissue exposure (AUC_0-last_) was observed in kidney (135000 ng·h/g) and lung (6840 ng·h/g), followed by small intestine, stomach, liver, large intestine, spleen and muscle. The heart, fat and brain exposures (AUC_0-last_) were not determined due to most concentrations were below the LLOQ (40 ng/g).

In regard to drug metabolism, FC-12738 was stable in CD-1 mouse, SD rat, beagle dog, cynomolgus monkey and human plasma after 120 minutes incubation and slowly metabolized in CD-1 mouse, SD rat, beagle dog, cynomolgus monkey and human hepatocytes. After incubation with plasma from CD-1 mouse, SD rat, beagle dog, cynomolgus monkey and human hepatocytes for 240 min, a total of 4 metabolites of FC-12738 were detected and identified. The metabolites were assigned as M1-M4: glucose conjugation metabolites (MW = 841.92, P + C6H10O5). Based on the identified metabolites, the major metabolic pathway of FC-12738 in hepatocytes was proposed as glucose conjugation. After incubation with human liver microsomes, no metabolites of FC-12738 were detected. Following a single IV dose of 2 mg/kg and SC dose of 10 mg/kg, 3 glucose conjugation metabolites (M2, M3, M4) were detected in rat plasma. Following single SC administration at the dose of 10 mg/kg FC-12738, 2 glucose conjugation metabolites (M3, M4) were detected in rat urine. No metabolites were detected in bile and feces.

In regard to drug excretion, following a single SC administration of FC-12738 at 10 mg/kg, the mean drug recovery of FC-12738 from urine and feces in the male intact rats at 168 hours post-dose were 86.7±6.25% and 2.19±1.22%, respectively, of the nominal dose. Drug recovery of FC-12738 from bile in the male bBDC rats at 72 hours post-dose was 0.336±0.260% of the nominal dose. These data indicate that urine was the main excretion route.

To examine potential drug-drug interactions we examined CYP450. The IC_50_ values of FC-12738 on CYP1A2, CYP2B6, CYP2C8, CYP2C9, CYP2C19, CYP2D6 and CYP3A (with midazolam and testosterone as probe substrate, respectively) were all greater than 100 μM and showed no time-dependent inhibition on any of the CYP isozymes tested. In the CYP450 induction study, FC-12738 at the concentrations ranging from 0.100 to 15.0 μM did not induce CYP1A2, CYP2B6 and CYP3A4. FC-12738 showed no inhibitory potential on the activity of efflux transports (P-gp and BCRP), the IC_50_ values of FC-12738 were both greater than 100 μM. Similarly, the IC_50_ values of FC-12738 were all greater than 100 μM for transporters OATP1B1/3, OAT1/3, OCT2, MATE1/2-K.

Collectively, these findings indicate that FC-12738 has improved stability over TP5 with low potential for adverse reactions. A study by Li et al reported that TP5 binds to TLR2 with a K_D_ of 1.57 μM, which equates to a fluid concentration of 1.060 ng/ml [1]. Assuming that FC-12738 has a similar affinity for TLR2, the levels of the peptide in brain were well above the K_D_ for binding the target for at least 2 hours post-injection. These findings suggest that FC-12738 could be utilized to manipulate neuroinflammation in neurodegenerative disease. Additional studies are required to determine the affinity of FC-12738 for TLR and its efficacy in target engagement *in vivo*.

## Acknowledgements

I thank Dr. David Borchelt PhD (University of Florida) for editorial assistance in preparing this report. Large portions of the text were provided by the contract research organization (WuXi App Tec Hong Kong Ltd) that conducted the study. The comparison study of TP5 and FC-12738 in CD-1 mice was conducted at the Fox Chase Chemical Diversity Center, Inc, which provided the description of methods used. The study was fully funded by Neurodegenerative Disease Research Inc. Dr. Ellison has filed a patent for FC-12738.

